# Proma Plasmids are Instrumental in the Dissemination of Linuron Catabolic Genes between Different Genera

**DOI:** 10.1101/831255

**Authors:** Johannes Werner, Eman Nour, Boyke Bunk, Cathrin Spröer, Kornelia Smalla, Dirk Springael, Başak Öztürk

## Abstract

PromA plasmids are broad host range plasmids, which are often cryptic and hence have an uncertain ecological role. We present three novel PromA γ plasmids which carry genes associated with degradation of the phenylurea herbicide linuron, two (pPBL-H3-2 and pBPS33-2) of which originate from unrelated *Hydrogenophaga* hosts isolated from different environments, and one (pEN1) which was exogenously captured from an on-farm biopurification system. Both *Hydrogenophaga* plasmids carry all three necessary gene clusters determining the three main steps for conversion of linuron to Krebs cycle intermediates, while pEN1 only determines the initial linuron hydrolysis step. Linuron catabolic gene clusters that determine the same step were identical on all plasmids, encompassed in differently arranged constellations and characterized by the presence of multiple IS*1071* elements. In all plasmids except pEN1, the insertion spot of the catabolic genes in the PromA γ plasmids was the same. Highly similar PromA plasmids carrying the linuron degrading gene cargo at the same insertion spot were were previously identified in linuron degrading *Variovorax* sp. Interestingly, in both *Hydrogenophaga* populations not every PromA plasmid copy carries catabolic genes. The results indicate that PromA plasmids are important vehicles of linuron catabolic gene dissemination, rather than being cryptic and only important for the mobilization of other plasmids.

## 1 INTRODUCTION

Plasmids are circular or linear extrachromosomal elements that can self-replicate, and are important agents in the dissemination of genes among microbial species (Garcillan-Barcia et al., 2011). Broad host range (BHR) plasmids can independently transfer and maintain themselves in different taxa (Jain and Srivastava, 2013) and carry accordingly genes for replication, maintenance and control, and conjugation (Szpirer et al., 1999). In addition, BHR plasmids may carry so-called “accessory” genes, for instance for antibiotic and heavy metal resistance, or biodegradation of xenobiotic compounds (Schluter et al., 2007; Sen et al., 2011). On the other hand, exogenous plasmid capture enabled the isolation of several plasmids with few or no apparent accessory genes that belong to known BHR plasmid groups such as IncP-1, IncN and IncU from environmental microbial communities (Brown et al., 2013). These so far cryptic plasmids have no apparent benefit to the host and are still propagated in absence of selective pressure (Fox et al., 2008). A recently discovered group of BHR plasmids are the PromA plasmids, most of which were isolated by exogenous plasmid capture (Thomas et al., 2017; Van der Auwera et al., 2009; Schneiker et al., 2001; Yanagiya et al., 2018; Li et al., 2014; Tauch et al., 2002) hence from unknown hosts, but also few derived from Proteobacterial isolates (Mela et al., 2008; Ito and Iizuka, 1971; Van der Auwera et al., 2009). With the exception of SFA231 (Li et al., 2014), pMOL98 (Van der Auwera et al., 2009) and pSB102 (Schneiker et al., 2001) which carry heavy metal resistance-related genes, all 12 completely sequenced PromA plasmids identified to date are cryptic plasmids with no clear indication of their ecological or possible benefit for the host organism. It was hypothesized that the main role of these plasmids is to mobilize other plasmids (Zhang et al., 2015).

Recently, we have described two PromA plasmids, pPBL-H6-2 and pPBS-H4-2 from two *Variovorax* strains with the metabolic capability to degrade the phenylurea herbicide linuron (Öztürk et al., 2019). *Variovorax* is a species that is isolated at high frequency from enrichments aiming for linuron degrading microorganisms. In the linuon-degrading *Variovorax* species isolated to date, the initial step of linuron degradation to 3,4-dichloroaniline (DCA) is performed by the linuron amidases *hylA* or *libA*, followed by the conversion of DCA to 4,5-dichlorocatechol by the *dcaQA1A2BR* catabolic cluster. The catechol intermediate is further degraded to Krebs cycle intermediates by the enzymes encoded by the *ccdCFDER* gene cluster (Bers et al. 2011, 2013). The two *Variovorax* PromA plasmids belong to the PromA γ subgroup together with the plasmids pSN1104-11 and pSN1104-34 (Yanagiya et al., 2018) that were exogeneously isolated from cow manure. pPBS-H4-2 is the first PromA plasmid that carries catabolic genes, i.e., it carries a stretch of DNA containing several gene clusters involved in the degradation of linuron and several IS*1071* elements, of which two border the cargo at both ends. In contrast, pPBL-H6-2 only contains one IS*1071* transposase as cargo.

In this study, we report on the characterization of three other linuron catabolic PromA γ plasmids. In contrast to pPBL-H6-2 and pPBS-H4-2, these plasmids did not originate from *Variovorax*. Two of the new plasmids were isolated from two different linuron degrading *Hydrogenophaga* strains, PBL-H3 and BPS33*. Hydrogenophaga* sp. strain PBL-H3 was isolated from a potato field near Halen, Belgium (Breugelmans et al., 2007) and *Hydrogenophaga* sp. strain BPS33 from the matrix of an on farm biopurification system (BPS) operated by Inagro, near Roeselare, Belgium, in this study. *Hydrogenophaga* is not a genus that is frequently associated with xenobiotic degradation. Exceptions comprise *Hydrogenophaga intermedia* S1 and PMC, which mineralize 4-aminobenzenesulfonate in two-species consortia (Gan et al., 2011), pyrene-degrading *Hydrogenophaga* sp. PYR1 (Yan et al., 2017) and 3-/4-hydroxybenzoate-degrading *Hydrogenophaga* sp. H7 (Fan et al., 2019). The third new PromA plasmid is plasmid pEN1, which was obtained by biparental exogenous isolation from a BPS operating on a farm near Kortrijk, Belgium (Dealtry et al., 2016), selecting for mercury resistant exconjugants of the recipient strain. We compared the full sequences of three new plasmids with each other and with other previously reported PromA plasmids including those discovered in the linuron degrading *Variovorax* in order to deduce their evolution and the role that PromA plasmids play in the dissemination of linuron degradation genes in different genera and environments.

## 2 MATERIALS AND METHODS

### 2.1 Chemicals

Linuron ([3-(3,4-dichlorophenyl)-1-methoxy-1-methyl urea] PESTANAL®, analytical standard) was purchased from Sigma Aldrich. [phenyl-U-14C] Linuron (16.93 mCi mmol^−1^, radio-chemical purity > 95%) was purchased from Izotop.

### 2.2 Isolation of *Hydrogenophaga* sp. BPS33

BPS33 was isolated from the matrix of a BPS located on the property of the research institute Inagro in Rumbeke-Beitem, Belgium (50°54’07.9”N 3°07’28.2”E). The BPS had received linuron and other pesticides for two years. The sample was collected from the upper 10 cm of the top container, and stored at 4°C until further use. The isolation procedure followed the protocol previously described by Breugelmans et al. (2007). Briefly, 1 g of the matrix material was inoculated into 50 ml MMO medium (Dejonghe et al., 2003) containing 20 mg/L linuron. Degradation of linuron was monitored using HPLC as described before (Horemans et al., 2014). After linuron was degraded, dilutions of the enrichment culture were plated on R2A medium (Breugelmans et al., 2007) containing 20 mg/L linuron. Resulting colonies were inoculated into 2.5 ml 96-well plates containing 500 µL of MMO with 20 mg/L linuron, and colonies that degraded linuron were identified via 16S rRNA gene sequencing with primers 27F and 1492R (Primers listed in Table S2). Both BPS33 and PBL-H3 used in this study were routinely cultivated in R2B supplemented with 20 mg/L linuron. Prior to genome sequencing, the mineralization capacity of both cultures was determined as described before (Breugelmans et al., 2007). Each mineralization test contained 10^8^ colony forming units of BPS33 or PBL-H3 in 40 ml MMO and a total radioactivity of 0.009 mCi mL^−1^.

### 2.3 Isolation of pEN1 by exogenous capture

Plasmid pEN1 was exogenously captured from a biopurification system material (Kortrijk, Belgium) spiked with linuron in a microcosm experiment (Dealtry et al., 2016). In brief, *Pseudomonas putida* KT2442 :*gfp* was used as a recipient strain for plasmids conferring mercury chloride resistance (20 µg mL-1). Biparental mating was performed with a bacterial suspension extracted from the matrix 25 day after linuron spiking.

### 2.4 Genome sequencing

DNA was isolated using Qiagen Genomic-tip 100/G (Qiagen, Hilden Germany) according to the instructions of the manufacturer. SMRTbell™ template library was prepared according to the instructions from PacificBiosciences, Menlo Park, CA, USA, following the Procedure & Checklist – Greater Than 10 kbp Template Preparation. Briefly, for preparation of 15 kbp libraries 8 µg genomic DNA and 1.4 µg plasmid DNA was sheared using g-tubes™ from Covaris, Woburn, MA, USA, according to the manufactureŕs instructions. DNA was end-repaired and ligated overnight to hairpin adapters applying components from the DNA/Polymerase Binding Kit P6 from Pacific Biosciences, Menlo Park, CA, USA. Reactions were carried out according to the manufactureŕs instructions. For the bacterial DNAs BluePippin™ Size-Selection to greater than 4 kbp was performed according to the manufactureŕs instructions (Sage Science, Beverly, MA, USA). Conditions for annealing of sequencing primers and binding of polymerase to purified SMRTbell™ template were assessed with the Calculator in RS Remote, PacificBiosciences, Menlo Park, CA, USA. 1 SMRT cell was sequenced per strain/plasmid on the PacBio RSII (PacificBiosciences, Menlo Park, CA, USA) taking one 240-minutes movies.

For the bacterial DNA libraries for sequencing on Illumina platform were prepared applying Nextera XT DNA Library Preparation Kit (Illumina, San Diego, USA) with small modifications (Baym et al., 2015). Samples were sequenced on NextSeq™ 500.

Bacterial long read genome assemblies were performed applying the RS_HGAP_Assembly.3 protocol included in SMRT Portal version 2.3.0 applying target genome sizes of 10 Mbp. For BPS33, the genome assembly directly revealed the chromosomal and both plasmid sequences. In case of PBL-H3, the assembly revealed the chromosomal sequence misassembled together with the 107 kbp plasmid. Thus, this plasmid sequence was separated from the chromosome and processed independently. Nevertheless, the 319 kbp plasmid was detected as separate contig. Further artificial contigs constituting of low coverage and/or included in other replicons were removed from the assembly. All remaining contigs were circularized; particularly assembly redundancies at the ends of the contigs were removed. Replicons were adjusted to *dnaA* (chromosome) or *repA-parA* (all plasmids) as the first gene. Error-correction was performed by a mapping of the Illumina short reads onto finished genomes using bwa v. 0.6.2 in paired-end (sampe) mode using default setting (Li and Durbin, 2009) with subsequent variant and consensus calling using VarScan v. 2.3.6 (Parameters: mpileup2cns –min-coverage 10 –min-reads2 6 –min-avg-qual 20 –min-var-freq 0.8 –min-freq-for-hom 0.75 –p-value 0.01 –strand-filter 1 –variants 1 –output-vcf 1) (Koboldt et al., 2012). A consensus concordance of QV60 could be reached. Automated genome annotation was carried out using Prokka v. 1.8 (Seemann, 2014). The hylA-containing plasmid was assembled using a target genome size of 200 kbp. However, only 25 percent of the plasmid population was shown to carry the transposon-based insertion based on plasmid coverage analysis coverage.

### 2.5 Assembly of pPBL-H3-2 variants B2 and B4

The pPBL-H3-2 variants were assembled using the pBPS33-2 as a scaffold. The PBL-H3 Illumina paired-end reads were mapped to pBPS33-2 using BWA-MEM v. 0.7.17.1 with standard settings (Li and Durbin, 2009) implemented in the Galaxy platform (Afgan et al., 2016). A consensus of the *dca* cluster genes together with the flanking regions was generated using samtools v. 2.1.4 mpileup with standard settings (Li et al., 2009). This consensus was aligned to pPBL-H3-2 (B2). The flanking regions matched pPBL-H3-2 (B2) perfectly, with *hylA* and the associated genes being located between these flanking sequences instead of the *dca* cluster. The consensus sequence with the *dca* cluster, acquired from the mpileup was inserted in the place of the *hylA* cluster in pPBL-H3-2 (B2) to obtain pPBL-H3-2 (B4). The new plasmid was annotated as described before. The assembly procedure is illustrated in Figure S2.

### 2.6 Comparative genomics analysis

Both phylogenetic trees and dDDH values were computed on the Type (Strain) Genome Server (TYGS) (Meier-Kolthoff and Goker, 2019). In brief, the TYGS analysis was subdivided into the following steps: The 16S rRNA gene sequences were extracted from the genomes using RNAmmer (Lagesen et al., 2007). All pairwise comparisons among the set of genomes were conducted using GBDP and intergenomic distances inferred under the algorithm ‘trimming’ and distance formula d_5_ (Meier-Kolthoff et al., 2013). Hundred distance replicates were calculated. Digital DDH values and confidence intervals were calculated using the recommended settings of the GGDC 2.1 (Meier-Kolthoff et al., 2013). The resulting intergenomic distances were used to infer a balanced minimum evolution tree with branch support via FASTME 2.1.4 including SPR postprocessing (Lefort et al., 2015). Branch support was inferred from 100 pseudo-bootstrap replicates each. The trees were rooted at the midpoint and visualized with iTOL (Letunic and Bork, 2019). For constructing the phylogenetic trees, six type strains and six additional *Hydrogenophaga* genomes were used to determine the phylogenetic position of the two linuron-degrading *Hydrogenophaga* strains within the genus. A dDDH species cutoff of 70% was applied as described before (Liu et al., 2015). The list of genomes included in the study is given in Supplementary table S1.

The alignment of the PromA plasmids was performed with AliTV v. 1.0.6 (Ankenbrand et al., 2017). Codon usage frequencies were calculated with Comparem v 0.0.23 (Parks, 2018), the PCA with R v. 3.5.2 (R Core Team, 2019) and FactoMineR v. 1.41 (Lê et al., 2008).

The plasmid sequences were categorized under known plasmid groups based on the aa and nucleotide identity of their backbone genes to known plasmids, using BLAST against the NCBI nr database (Altschul et al., 1990). To elucidate the RepA gene-based phylogeny of the PromA plasmids, nucleotide sequences were aligned with MUSCLE v. 3.8.31 (Madeira et al., 2019; Edgar, 2004) and the maximum likelihood (ML) trees were calculated with RaxML v. 8 (Stamatakis, 2014) using the under the GTR+GAMMA model and 1000 bootstrap replicates. The genomic locations of the IS*1071* elements and catabolic genes were determined and the genes were visualized using Geneious v. 11.0.4.

### 2.7 Quantification of bacteria, plasmids and catabolicgenesby real-time quantitative PCR (qPCR)

The PCR-qPCR primer sequences used in this study are listed in Supplementary table S2. For the qPCR analysis, the strains BPS33 and PBL-H3 were grown to OD_600_ as described above. DNA extraction was made from 2 ml of culture as previously-described (Larsen et al., 2007). Each reaction contained 10 ng of template DNA. qPCR reactions were performed with the ABsolute QPCR Mix (Thermo Fisher) on a Roche LightCycler 480 II. The qPCRs for 16S rRNA gene (Lopez-Gutierrez et al., 2004), *hylA* (Horemans et al., 2016) and *dcaQ* (Albers et al., 2018) quantification were performed as previously described. qPCR targeting korB was performed as previously-described, except that the Taqman probe was omitted (Jechalke et al., 2013). Each PCR reaction to generate the templates for the qPCR standard curves contained 10 ng template DNA (BPS33 gDNA), 1x Dream Taq Green buffer (Thermo Fisher), 0.2 M of each dNTP, 0.1 µM of each primer and 1.25 U of Dream Taq DNA polymerase (Thermo Fisher) in a final volume of 50 µl. The amplification was performed as follows: Initial denaturation of 95°C for 3 min, 40 cycles of denaturation at 95°C for 30 s, annealing at 60°C for 30 s, extension at 72°C for 1 min, followed by a final extension at 72°C for 15 min. All conventional PCR reactions were performed with an Applied Biosystems Veriti 96-well thermal cycler. The products were purified with the DNA Clean&Concentrator 25 kit (Zymo) and quantified with the Qubit BR DNA assay (Thermo Fisher).

## 3 RESULTS

### 3.1 Phylogenetic analysis of the PromA plasmids

*Hydrogenophaga* plasmids pPBL-H3-2, pBPS33-2 and exogenously-captured pEN1, were completely sequenced. The general features of plasmids pEN1, pPBL-H3-2 and pBPS-33 are given in Table 1.

**Table 1.** Statistics of the genomes and plasmids sequenced in this study.

|  | contig<br>size | #CDS | %GC | # rRNA<br>operons | #tRNA | classification |
| --- | --- | --- | --- | --- | --- | --- |
| pEN1 | 59 kbp | 68 | 63.4 |  |  | PromA plasmid |
| chr BPS33 | 6.33 Mbp | 5,830 | 65.7 | 2 | 53 | chromosome |
| pBPS33-1 | 340 kbp | 323 | 61.2 |  |  | unclassified<br>plasmid |
| pBPS33-2 | 107 kbp | 109 | 63.1 |  |  | PromA plasmid |
| chr PBL-H3 | 4.39 Mbp | 4.39 | 65.6 | 2 | 44 | chromosome |
| pPBL-H3-1 | 320 kbp | 320 | 61.1 |  |  | unclassified<br>plasmid |
| pPBL-H3-2 | 107 kbp | 107 | 62.8 |  | 1 | PromA plasmid |

A whole-sequence alignment revealed that the plasmids pEN1, pBPS-33-2 and pPBL-H3-2 are very closely related to the previously-described plasmids pPBS-H4-2 and pPBL-H6-2 from linuron-degrading *Variovorax* sp. (Öztürk et al., 2019) (Figure 1). Indeed, RepA-based phylogenetic analysis showed that these plasmids all belong to the PromA γ group, together with the plasmids pSN1104-11 and pSN1104-34 (Yanagiya et al., 2018) (Figure 2).

**FIGURE 1.**
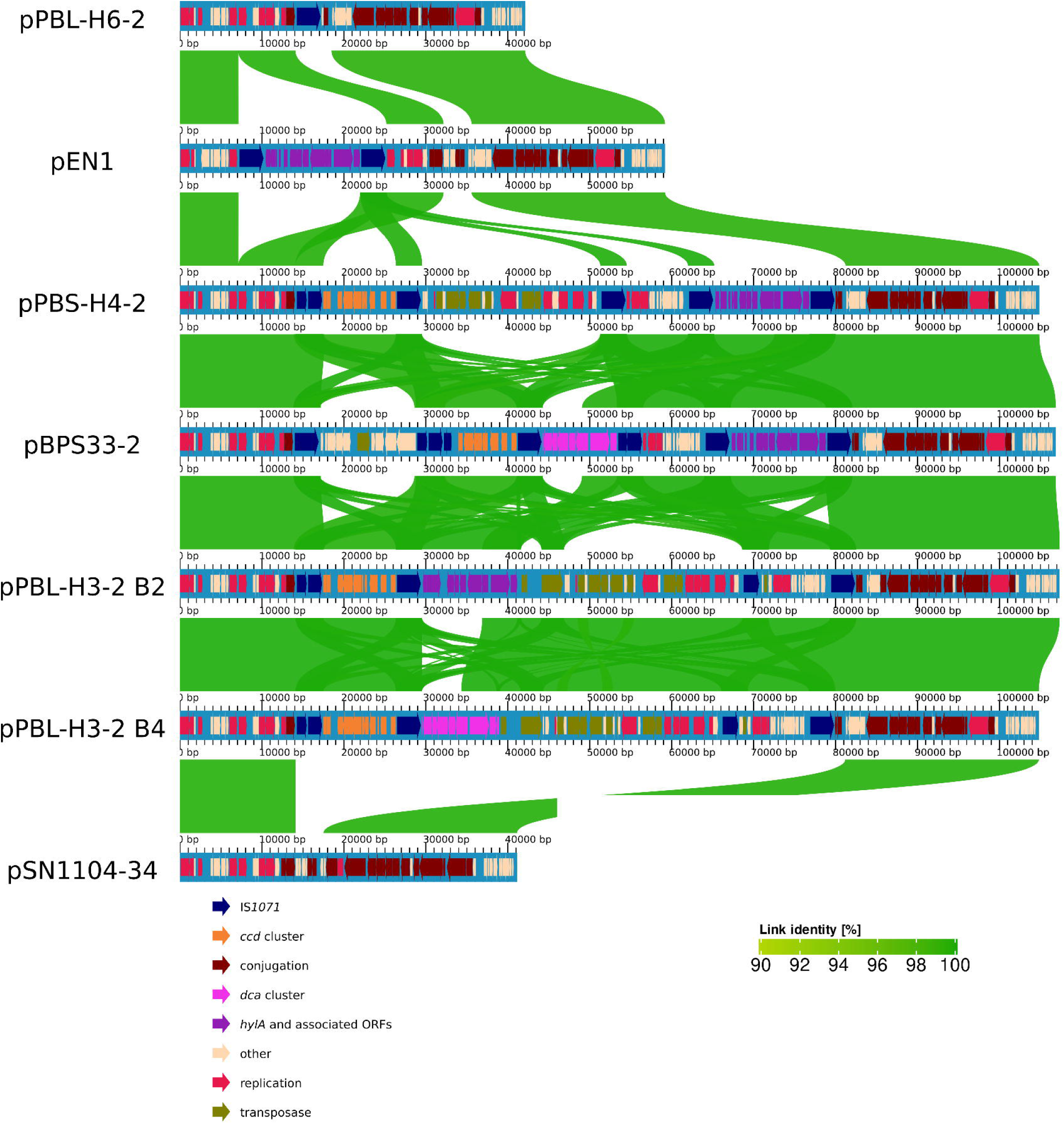
Alignment of PromA γ plasmids. Alignment identities are shown for an identity of 90-100%. Each plasmid is aligned to the one above.

**FIGURE 2.**
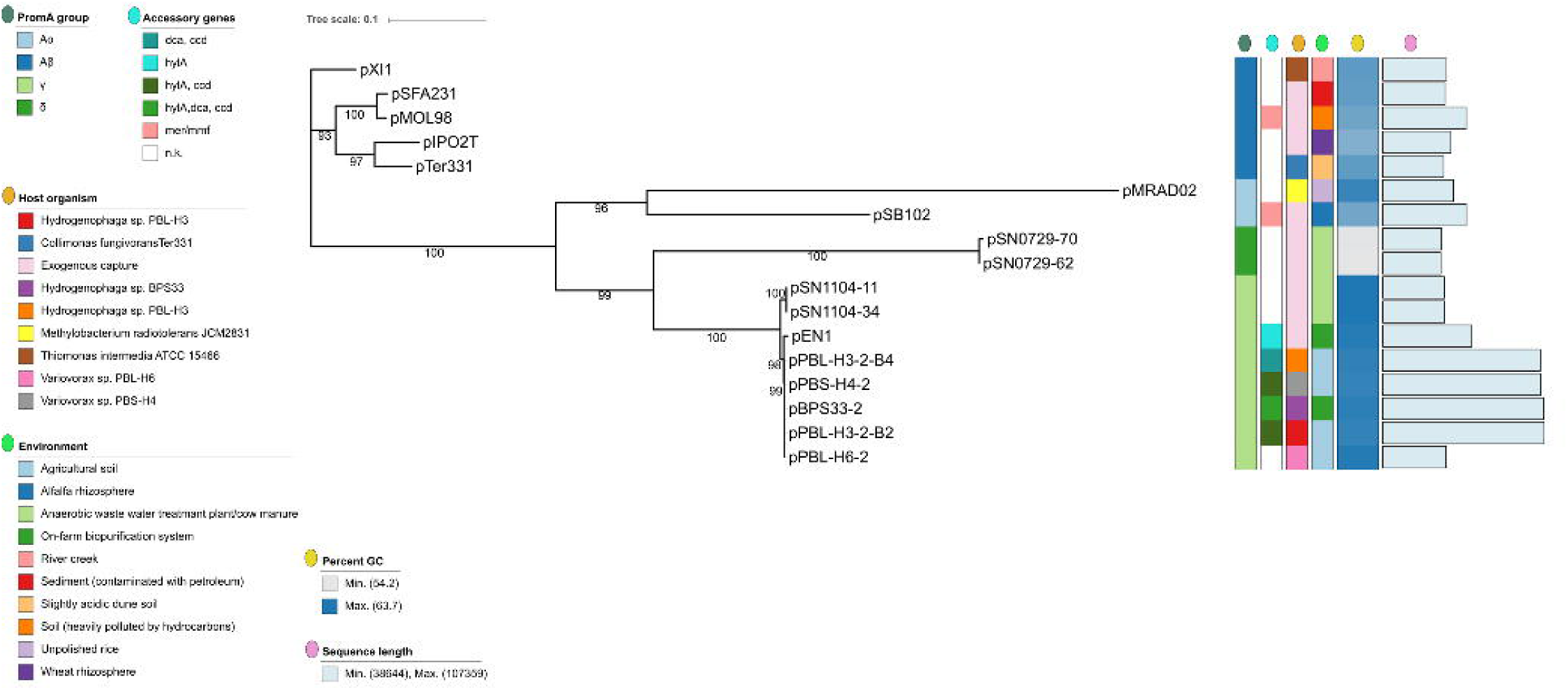
*repA* gene-based phylogeny of PromA plasmids. The branches are scaled in terms of the expected number of substitutions per site. The numbers above the branches are support values when larger than 60 % from ML.

The RepA sequences, as well as the type IV secretion system sequences of the PromA γ plasmids were highly-conserved, with 99% identity to each other on amino acid level. Codon usage-based clustering showed that catabolic PromA γ plasmids clustered together and separately from the non-catabolic ones (Figure 3).

**FIGURE 3.**
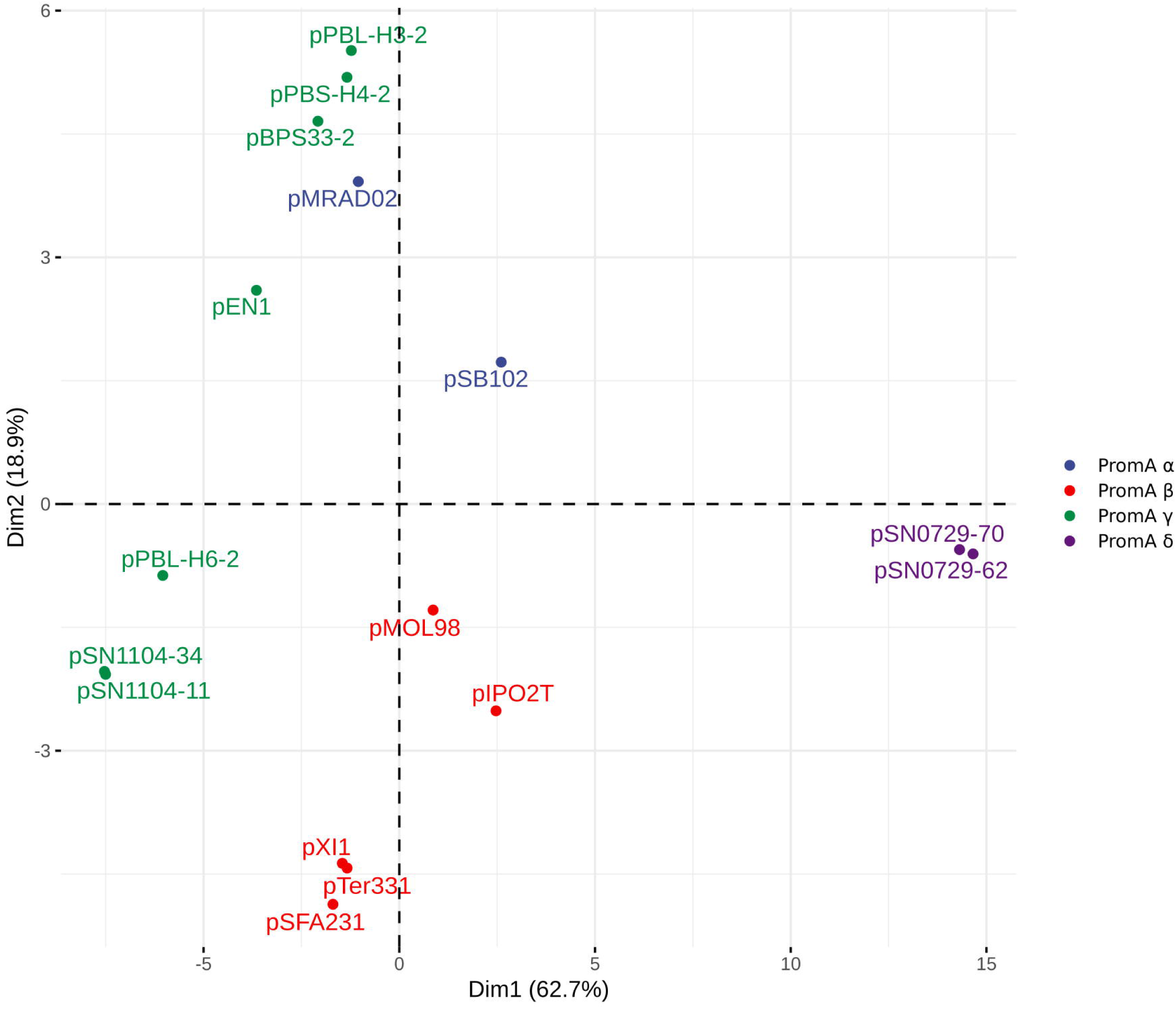
PCA of the codon usage frequencies of the promA plasmids. PromA δ plasmids are separated by the first dimension from the other plasmids, and PromA β plasmids by the second dimension

### 3.2 Catabolic potential of the PromA plasmids

The newly-sequenced PromA plasmids pBPS33-2, pPBL-H3-2 and pEN1 all carry genes related to linuron-degradation. After the assembly of the PBL-H3 genome, it was observed that the genome did not contain the *dca* cluster, which was not expected due to the capacity of this strain to mineralize linuron completely. A BLAST search against the unassembled PacBio reads indeed revealed the presence of the same *dca* cluster genes as those present on pBPS33-2. The new assembly, which makes use of the similarity between the two plasmids as described in the methods section, revealed that PBL-H3 had two different versions of the pPBL-H3-2 within the population, named as pPBL-H3-2 (B2) and pPBL-H3-2 (B4). Both pPBL-H3-2 (B2) and pPBL-H3-2 (B4) are identical except that in the catabolic gene cluster the locus carrying *hylA* gene and associated open reading frames (ORFs), is replaced by the *dca* cluster gene cluster (Figure 1).

pBPS33-2 carries all the genes necessary for linuron degradation, while pEN1 only contains *hylA*. *hylA*, *dcaQA1A2BR* and *ccdCFDER* are 99% identical to those previously identified in *Variovorax* sp. WDL1 and PBS-H4. The *hylA* gene on all plasmids truncates a *dcaQ* gene, with the junction being identical to the *hylA*-*dcaQ* junction in pPBS-H4-2 (Öztürk et al., 2019). pEN1 on the other hand carries a *hylA*-*dcaQ* junction identical to that on pWDL1-1, the *Variovorax* sp. WDL1 megaplasmid that carries the linuron degradation genes of this bacterium (Albers et al., 2018; Öztürk et al., 2019). As previously reported for pWDL1-1 and pPBS-H4-2, the catabolic clusters on the newly-sequenced PromA plasmids are flanked by IS*1071*-class II insertion elements, forming putative composite transposons. In pBPS33-2, the *ccd* and *dca* clusters are adjacent to each other, with one IS*1071* in between, amounting to three IS elements in total. The IS*1071* element associated with *hylA* is separated from the *dca* and *ccd* and flanked by two additional IS*1071* elements.

In addition to the catabolic clusters related to linuron degradation, pBPS33-2 carries an extra IS*1071* element flanking genes that encode for four proteins putatively involved in the *meta*-pathway of phenol degradation and three putative multidrug efflux pump proteins, with an intermittent a single copy of an IS*91*-class transposase, encompassing in total 18 kbp.

### 3.3 PromA plasmid associated IS*1071* insertion elements and their synteny

The catabolic PromA plasmids have a high number of IS*1071*-elements, some of which are associated with the catabolic clusters. IS*1071* elements are absent in the non-catabolic PromA plasmids, with the exception of pPBL-H6-2 (Öztürk et al., 2019). The IS*1071* element sequences are largely similar to the classical structure (Sota et al., 2006); except that some elements seem to code for truncated transposase due to a premature stop codon (Figure 4). pBPS33-2 has six IS*1071* elements, the highest number among all PromA plasmids described so far, followed by five in pPBS-H4-2, four in pPBL-H3-2, two in pEN1 and one in pPBL-H6-2.

**FIGURE 4.**
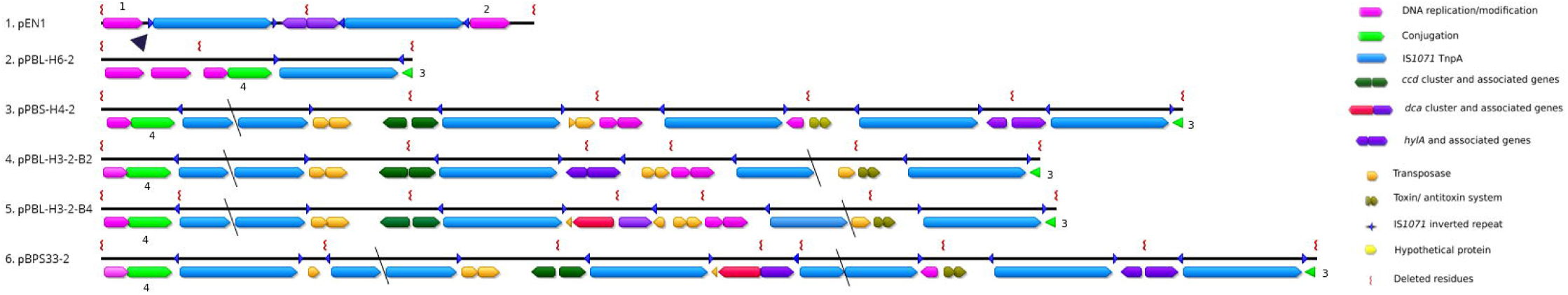
The IS*1071* insertion sites on various PromA γplasmids. The residues between the IS*1071* transposon were deleted for simplicity, except the immediate flanking genes. The insertion locus of the pEN1 IS*1071* was marked on pPBL-H6-2 with a triangle, and is conserved in other PromA-γ plasmids. The slanted black lines on the IS*1071* ORF indicate immature stop codons. The IS*1071*-flanking backbone genes are numbered 1-4: (1) *repB* (2) *mobA* (3) *trbM* (4) *VirD2*.

The IS*1071* insertion sites among the plasmids pBPS33-2, pPBL-H3-2 and pPBS-H4-2 are highly conserved, the first insertion site relative to *repA* being adjacent to the plasmid mobility genes *mobC* and *virD2* and the 3’ end of the last IS*1071* element being flanked by the backbone *trbM* gene (Figure 4). pPBL-H6-2 only has one IS*1071* transposase with inverted repeats (IR). For pBPS33-2, pPBL-H3-2 (B2/B4) and pPBS-H4-2, the first insertion site has been subject to multiple transposon insertion events, where multiple catabolic clusters and other accessory genes have been inserted consecutively, being only separated by one IS*1071* element including IRs. Interestingly, some genes that are associated with DNA replication, such as the genes encoding for the plasmid replication and segregation proteins RepA and RepB (100% aa identity to WP_068682750.1 and WP_068682748.1, respectively), segregation proteins ParA and ParB (100% aa identity to CDS81791.1 and WP_011114060.1, respectively), tyrosine recombinase XerC (100% aa identity to WP_068682758.1) and toxin/antitoxin system genes *klcA/dinJ/yafQ* (100% aa identity to WP_011114069.1, WP_011114070.1 and WP_011114071.1, respectively), are also flanked by IS*1071* elements. pPBS-H4-2 and pPBL-H3-2 (B2/B4) carry eight DNA replication-related genes flanked by IS*1071* elements, which are 100% identical to each other on nucleotide level, while pBPS33-2 carries three of these eight genes, which are identical to their pPBS-H4-2/pPBL-H3-2 counterparts (*parAB and kfrA*). The other PromA γ plasmids lack the IS*1071*-associated DNA-replication related genes as well as the toxin/antitoxin system genes altogether. On the other hand, all PromA γ plasmids have copies of *parA* and *parB* genes, which are unrelated to those flanked by the IS*1071* elements on the catabolic plasmids. In addition to these accessory genes, six putative transposases and thirteen hypothetical proteins were found in the IS*1071*-flanked region in pPBS-H4-2 and pPBL-H3-2 (B2/B4), which are conserved among those two. Six of them are present in pBPS33-2, all of which are annotated as hypothetical proteins. These are absent in the other PromA γ plasmids.

The insertion site of the pEN1 IS*1071* element is an exception among the PromA plasmids. The insertion site of this IS*1071* element, associated with the *hylA* gene and adjacent ORFs (Albers et al., 2018; Öztürk et al., 2019), lies between the backbone genes *repB* and *mobA*. This site is located before the first insertion site of the other catabolic plasmids relative to *repA*, and exists in other PromA γ plasmids, but with four nucleotide differences. The *repB-mobA* intergenic region on plasmid SN1104-34, which has no IS*1071* elements, is identical to pPBS-H4-2, pPBL-H3-2 and pBPS-33-2, except for a single nucleotide. The left IR of the first IS1071 element on pEN1 has twelve nucleotide differences to the previously-described IS*1071* left IR (Sota et al., 2006). The left IR of the second IS*1071* on pEN1, just like all the other left IRs on the other catabolic PromA plasmids, is identical to what was previously described (Sota et al., 2007).

In pPBL-H3-2 (B2/B4), the IS*1071* element flanking the right side of the *hylA/dca* catabolic cluster appears to encode a truncated variant of the IS*1071* transposase, which is not the case in pBPS33-2 and pPBS-H4-2, where both IS*1071* transposases are intact (Figure 4). On PBL-H3-2 (B2/B4) and pBPS33-2 the *ccd* cluster is adjacent to the *hylA*/*dca* clusters, with one IS*1071* element in between. The left flanking IS*1071* transposase of the *ccd* cluster appears truncated in all PromA plasmids at identical positions. In all cases, truncations were caused by an immature stop codon as a result of a point mutation. This truncated transposase was not present in the pWDL1 of *Variovorax* sp. WDL1, where an identical *ccd* cluster to those on pPBL-H3-2 (B2 and B4), pBPS33-2 and pPBS-H4-2 is located.

### 3.4 PromA plasmid and catabolic gene copy numbers in *Hydrogenophaga*

As there were two variants of the pPBL-H3-2 present in the PBL-H3 population, the question arose whether each version is represented in equal numbers, and how this compares to the BPS33 population where only one plasmid version could be assembled. The copy numbers of the genes encoding for KorB, HylA, DcaQ, as well as 16S rRNA were determined to elucidate both the number of PromA plasmid copies (*korB*) per cell and the proportions of PromA plasmids that carry the catabolic genes. Cultures of PBL-H3 and BPS33 contained approximately 10 copies of the *korB* gene per cell, hence 10 copies of the PromA plasmid per cell. The *hylA* and *dcaQ* gene copy numbers in both strains were similar, at about one copy per 100 cells, i.e., about one copy per 1000 PromA plasmid copies.

### 3.5 General genome characteristics and phylogeny

The general genome characteristics of the two *Hydrogenophaga* strains are summarized in Table 1. Prior to sequencing, the ability of both strains to mineralize linuron was confirmed as described in the Materials and Methods section. Both genomes comprise of one chromosome and two plasmids.

To the date of this publication, 22 *Hydrogenophaga* genomes have been deposited to GenBank, five of which are complete genomes. The complete list of genomes included in the phylogenetic analysis is given in Table S1. The complete genomes have a size of 4.39-6.32 Mbp, with GC contents ranging from 61 to 70%. BPS-33 has the largest genome of them with 6.32 Mbp. This is an exceptional size, the nearest largest *Hydrogenophaga* genome being 5.23 Mbp. Apart from the linuron-degrading strains sequenced in this study, only *Hydrogenophaga pseudoflava* DSM 1084 (pDSM1084, NZ_CP037868.1, 45.2 kbp) contains plasmids.

Phylogenetic analysis revealed a distant relation of PBL-H3 and BPS33 to each other (Figure 5). The computed DNA-DNA hybridization of 25.7% confirms that these two isolates are different species. The isolates do not belong to any type species.

**FIGURE 5.**
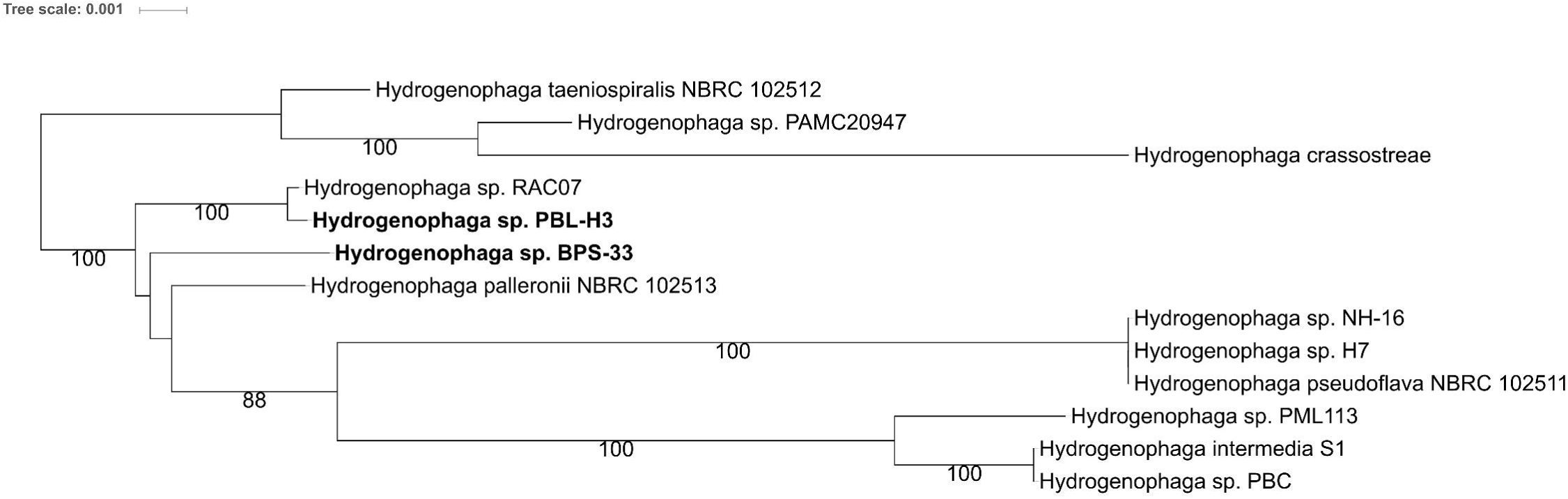
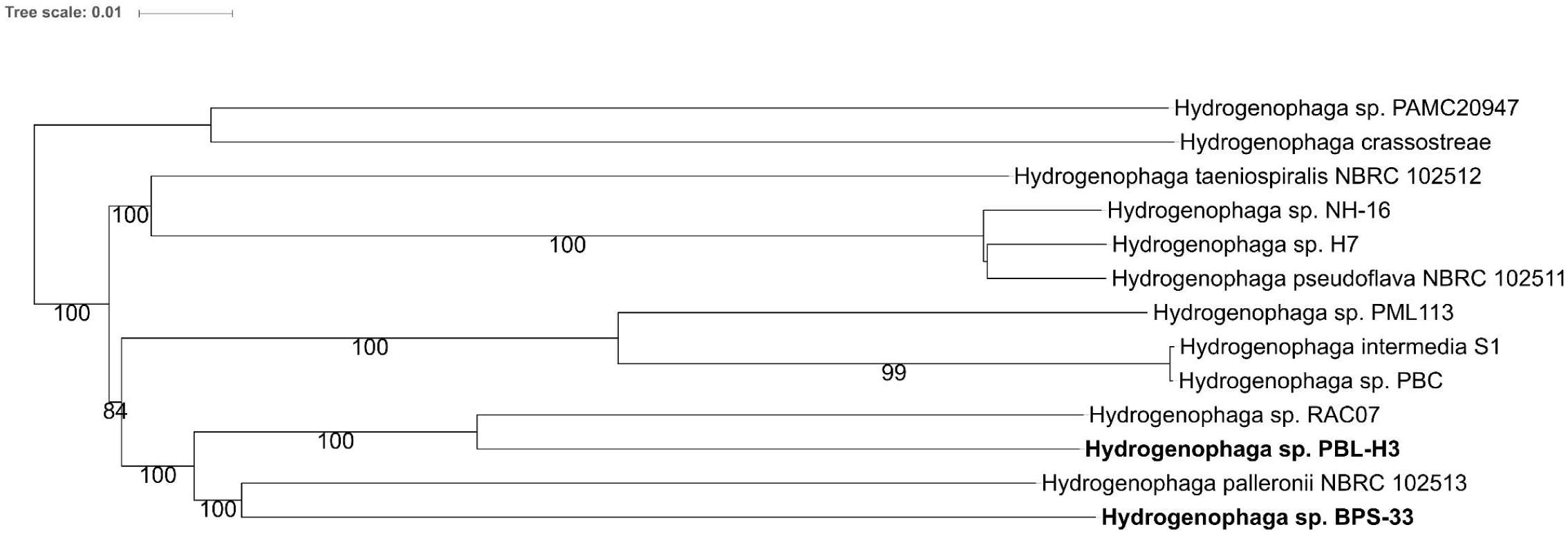
Phylogeny of Hydrogenophaga species with fully-sequenced based on genomes on (a) whole genome sequences and (b) 16S rRNA gene sequences. The linuron-degrading Hydrogenophaga sp. PBL-H3 and BPS33 sequenced in this study are marked in bold. The branch lengths are scaled in terms of GBDP distance formula d5. The numbers above branches are GBDP pseudo-bootstrap support values > 60% from 100 replications.

Both *Hydrogenophaga* strains carried megaplasmids, pBPS33-1 and pPBL-H3-1, which could not be assigned to any known plasmid class. The plasmids do not share any similarity and have no similarity to megaplasmids previously identified in *Variovorax* (Öztürk et al., 2019) or any other plasmid from *Hydrogenophaga*.

## 4 DISCUSSION

In this study, we have investigated the genomic basis of linuron degradation by the two *Hydrogenophaga* strains PBL-H3 and BPS33, and the role of PromA plasmids in the dissemination of catabolic genes in different environments. The genomes of two linuron-degrading *Hydrogenophaga* strains were completely sequenced and chromosomes and plasmids were circularized. These strains were sampled from two different environments, and although they belong to the same genus, they are distantly related to each other. The linuron-degrading *Variovorax* strains were very closely related to each other despite belonging to different species (Öztürk et al., 2019). Within the *Hydrogenophaga* genus however, the ability to acquire xenobiotic degradation genes seems to be independent of the host phylogeny.

The PromA plasmids pPBL-H3-2 and pBPS33-2 are the sole carriers of linuron-related catabolic clusters in both strains analyzed in this study. This was also the case for *Variovorax* sp. PBS-H4, although this strain lacks the *dca* cluster genes that are required for complete linuron mineralization (Öztürk et al., 2019). Remarkably, the catabolic PromA plasmids have a near-identical backbone to the previously-described PromA γ plasmids isolated in Japan (Yanagiya et al., 2018). It has been reported before that BHR plasmids isolated from different geographic locations can be highly-conserved (Li et al., 2016; Heuer et al., 2004; Chen et al., 2015b), however, some degree of divergence in the backbone structures of PromA plasmid groups were reported before (Li et al., 2014). In this case, the near-identical backbone genes indicate that these plasmids had a recent common ancestor, probably without any accessory genes. Interestingly, the PromA plasmids of PBL-H3 (B2) and PBS-H4, which were isolated from the same soil sample, have a high synteny of their cargo genes, indicating that these plasmids are variants of each other and can possibly be transferred within these two genera in the same environment. The codon usage distribution of the closely related PromA γ plasmids differ among those with and without catabolic clusters, indicating that the cargo genes differ in codon usage from the backbone genes.

IS*1071* elements were associated with all catabolic clusters related to linuron degradation on all plasmids. Remarkably, all catabolic clusters were near-identical to what has previously been described for *Variovorax* (Öztürk et al., 2019), both in terms of gene identity and synteny. This demonstrates that even among different genera and environments, the linuron-degradation pathways rely on a limited genetic repertoire, and the role of IS*1071* elements to transfer these genes is not limited to a certain genus or environment. The cargo associated by IS*1071* elements on these plasmids, however, were not limited to catabolic genes. Putative plasmid backbone genes involved in plasmid replication and maintenance as well as hypothetical proteins were also associated with IS*1071* elements, which were absent in the PromA plasmids without IS*1071* elements. The nearest relatives of the plasmid backbone genes associated with IS*1071* elements originated from different organisms and plasmids, among which are IncP-1 plasmids (*parAB*) but also non-linuron-degrading *Variovorax* chromosomes (*repAB*). It is worth noting that IS*1071* type transposases were not the only transposases located on some of the PromA plasmids. Especially pPBS-H4-2 and pPBL-H3-2 (B2/B4) carry a number of different transposases, showing that PromA plasmids are prone to acquiring mobile genetic elements and driving horizontal gene transfer.

The identical IS*1071* insertion sites on the PromA γ plasmids indicate hot spots for transposon insertion. Hot spots were previously-reported for IncP plasmids with IS*1071* elements (Sota et al., 2007; Dunon et al., 2018; Thorsted et al., 1998). The insertion of transposons at specific sites contributes to plasmid stability (Sota et al., 2007). The hot spot on our PromA γ plasmids is located between the genes encoding for the relaxase VirD2 and conjugal transfer protein TrbM, which is different from the *parA* locus that was previously shown to be the insertion site of pSFA231 (Li et al., 2014), pMOL98 (Van der Auwera et al., 2009) and pSB102 (Schneiker et al., 2001) as well as using PCR-based methods in metagenomes (Dias et al., 2018). The PromA γ insertion hot spot is in some cases occupied by multiple consecutive IS elements. The counterparts of the insertion site on pEN1 on the other catabolic plasmids on the other hand, are not occupied despite the fact that these plasmids all carry a high number of IS*1071* elements.

The qPCR results indicate that the *hylA* and *dca* cluster carrying PromA plasmids are underrepresented by almost 100-fold compared to the total number of PromA plasmids in both species. This holds true for pPBL-H3-2, for which two variants carrying either *hylA* or *dca* genes were assembled, as well as pBPS33-2. The qPCR results show that both BPS33 and PBL-H3 populations harbour different PromA plasmids, of which not all have the *hylA* gene or the *dca* cluster, and the majority lack both. Isogenic subpopulations carrying either *hylA* or *dca* genes were reported for *Variovorax* sp. WDL1 before (Albers et al., 2018). It was proposed that the existence of two subpopulations may be an adaptation to linuron degradation in a consortium, where linuron is degraded to DCA *hylA*-carrying consortium member, while the DCA degradation is performed by the other consortium members (Albers et al., 2018). Indeed, PBL-H3 indeed tends to accumulate DCA when growing on linuron on its own, and performed much better in a consortium with other DCA degraders (Breugelmans et al., 2007). Thus, *Hydrogenophaga* strains might be adapted in a similar way. Interestingly, the linuron catabolic genes of both *Hydrogenophaga* strains are near-identical to those of *Variovorax* strains WDL1 and PBS-H4, where WDL1 degrades linuron less efficiently on its own than when it is a part of a consortium (Dejonghe et al. 2003), and PBS-H4 can only perform the conversion of linuron to DCA (Breugelmans et al. 2007). The major difference between the *Variovorax* sp. WDL1 and *Hydrogenophaga* subpopulations is that, in WDL1 the majority of the population has either the *hylA* gene or the *dca* cluster (Albers et al., 2018), while in both *Hydrogenophaga* strains subpopulations containing either gene are much underrepresented.

The results show that even among different genera, the genes for complete linuron mineralization are highly-conserved, being acquired through horizontal gene transfer which is mediated by BHR plasmids. PromA γ plasmids, in addition to the previously-known IncP-1 plasmids, are carriers of IS*1071* elements and associated catabolic pathways, being present in different contaminated ecosystems. In contrast to the linuron-degrading *Variovorax* species, where the degradation genes are also found on megaplasmids as well as BHR plasmids (Öztürk et al., 2019), the *Hydrogenophaga* catabolic genes are only found on BHR plasmids, pointing towards a more recent acquisition of these gene clusters.

### 4.1 Data availability

The BPS33 genome is available under the accession numbers CP044549-CP044551, PBL-H3 (B2) under CP044975-CP044977, PBL-H3 (B4) under CP044972-CP044974 and pEN1 plasmid sequence under MN536506.

## ACKNOWLEDGEMENTS

This work was supported by the EU 7th Framework Programme (MetaExplore 222625) and FWO Project G.0371.06. We thank Anja Heuer and Simone Severitt for technical assistance, Charlotte Roschka for her help in sequence analysis and assembly, and Jörg Overmann for his support for sequencing of the strains. Johannes Werner personally acknowledges the use of de.NBI cloud and the support by the High Performance and Cloud Computing Group at the Zentrum für Datenverarbeitung of the University of Tübingen and the Federal Ministry of Education and Research (BMBF) through grant no 031 A535A.

## AUTHOR CONTRIBUTIONS STATEMENT

BÖ designed the study and performed the experiments on the *Hydrogenophaga* strains. BÖ and JW performed the sequence analysis of the *Hydrogenophaga* genomes and comparative analysis of PromA plasmids. BB and CS performed the sequencing and assembly of all genomes and plasmids. EN and KS isolated pEN1 and performed the sequence analysis. BÖ, JW, and DS wrote the main body of the paper. All authors contributed to the writing and critical reading of this publication.

## REFERENCES

Afgan, E., Baker, D., van den Beek, M., Blankenberg, D., Bouvier, D., Cech, M., Chilton, J., Clements, D., Coraor, N., Eberhard, C., Gruning, B., Guerler, A., Hillman-Jackson, J., Von Kuster, G., Rasche, E., Soranzo, N., Turaga, N., Taylor, J., Nekrutenko, A. and Goecks, J. (2016) The Galaxy platform for accessible, reproducible and collaborative biomedical analyses: 2016 update. Nucleic Acids Res., 44, W3–W10.

Albers, P., Lood, C., Ozturk, B., Horemans, B., Lavigne, R., van Noort, V., De Mot, R., Marchal, K., Sanchez-Rodriguez, A. and Springael, D. (2018) Catabolic task division between two near-isogenic subpopulations co-existing in a herbicide-degrading bacterial consortium: consequences for the interspecies consortium metabolic model. Environ. Microbiol., 20, 85–96.

Altschul, S. F., Gish, W., Miller, W., Myers, E. W. and Lipman, D. J. (1990) Basic local alignment search tool. J Mol Biol, 215, 403–410.

Ankenbrand, M. J., Hohlfeld, S., Hackl, T. and Förster, F. (2017) AliTV—interactive visualization of whole genome comparisons. PeerJ Comp Sci, 3, e116.

Van der Auwera, G. A., Krol, J. E., Suzuki, H., Foster, B., Van Houdt, R., Brown, C. J., Mergeay, M. and Top, E. M. (2009) Plasmids captured in *C. metallidurans* CH34: defining the PromA family of broad-host-range plasmids. Antonie Van Leeuwenhoek, 96, 193–204.

Baym, M., Kryazhimskiy, S., Lieberman, T. D., Chung, H., Desai, M. M. and Kishony, R. (2015) Inexpensive multiplexed library preparation for megabase-sized genomes. PLoS ONE, 10, e0128036.

Bers K et al. 2011. A novel hydrolase identified by genomic-proteomic analysis of phenylurea herbicide mineralization by *Variovorax* sp. strain SRS16. Appl. Environ. Microbiol.

Bers K et al. 2013. Hyla, an alternative hydrolase for initiation of catabolism of the phenylurea herbicide linuron in *Variovorax* sp. strains. Appl. Environ. Microbiol. 79:5258–5263.

Breugelmans, P., D’Huys, P. J., De Mot, R. and Springael, D. (2007) Characterization of novel linuron-mineralizing bacterial consortia enriched from long-term linuron-treated agricultural soils. FEMS Microbiol. Ecol., 62, 374–385.

Brown, C. J., Sen, D., Yano, H., Bauer, M. L., Rogers, L. M., Van der Auwera, G. A. and Top, E. M. (2013) Diverse broad-host-range plasmids from freshwater carry few accessory genes. Appl. Environ. Microbiol., 79, 7684–7695.

Chen, J., Bhattacharjee, H. and Rosen, B. P. (2015a) ArsH is an organoarsenical oxidase that confers resistance to trivalent forms of the herbicide monosodium methylarsenate and the poultry growth promoter roxarsone. Mol. Microbiol., 96, 1042–1052.

Chen, K., Xu, X., Zhang, L., Gou, Z., Li, S., Freilich, S. and Jiang, J. (2015b) Comparison of four *Comamonas* catabolic plasmids reveals the Evolution of pBHB To catabolize haloaromatics. Appl. Environ. Microbiol., 82, 1401–1411.

Dealtry, S., Nour, E. H., Holmsgaard, P. N., Ding, G. C., Weichelt, V., Dunon, V., Heuer, H., Hansen, L. H., Sørensen, S. J., Springael, D. and Smalla, K. (2016) Exploring the complex response to linuron of bacterial communities from biopurification systems by means of cultivation-independent methods. FEMS Microbiol. Ecol., 92.

Dejonghe, W., Berteloot, E., Goris, J., Boon, N., Crul, K., Maertens, S., Hofte, M., De Vos, P., Verstraete, W. and Top, E. M. (2003) Synergistic degradation of linuron by a bacterial consortium and isolation of a single linuron-degrading *Variovorax* strain. Appl. Environ. Microbiol., 69, 1532–1541.

Di Gioia, D., Peel, M., Fava, F. and Wyndham, R. C. (1998) Structures of homologous composite transposons carrying *cbaABC* genes from Europe and North America. Appl. Environ. Microbiol., 64, 1940–1946.

Dias, A. C. F., Cotta, S. R., Andreote, F. D. and van Elsas, J. D. (2018) The *parA* region of broad-host-range PromA plasmids is a carrier of mobile genes. Microb. Ecol., 75, 479–486.

Dunon, V., Bers, K., Lavigne, R., Top, E. M. and Springael, D. (2018) Targeted metagenomics demonstrates the ecological role of IS*1071* in bacterial community adaptation to pesticide degradation. Environ. Microbiol., 20, 4091–4111.

Dunon, V., Sniegowski, K., Bers, K., Lavigne, R., Smalla, K. and Springael, D. (2013) High prevalence of IncP-1 plasmids and IS*1071* insertion sequences in on-farm biopurification systems and other pesticide-polluted environments. FEMS Microbiol. Ecol., 86, 415–431.

Edgar, R. C. (2004) MUSCLE: multiple sequence alignment with high accuracy and high throughput. Nucleic Acids Res., 32, 1792–1797.

Fan, X., Nie, L., Shi, K., Wang, Q., Xia, X. and Wang, G. (2019) Simultaneous 3-/4-hydroxybenzoates biodegradation and arsenite oxidation by *Hydrogenophaga* sp. H7. Front Microbiol, 10, 1346.

Fox, R. E., Zhong, X., Krone, S. M. and Top, E. M. (2008) Spatial structure and nutrients promote invasion of IncP-1 plasmids in bacterial populations. ISME J, 2, 1024–1039.

Gan, H. M., Shahir, S., Ibrahim, Z. and Yahya, A. (2011) Biodegradation of 4-aminobenzenesulfonate by *Ralstonia* sp. PBA and Hydrogenophaga sp. PBC isolated from textile wastewater treatment plant. Chemosphere, 82, 507–513.

Garcillan-Barcia, M. P., Alvarado, A. and de la Cruz, F. (2011) Identification of bacterial plasmids based on mobility and plasmid population biology. FEMS Microbiol. Rev., 35, 936–956.

Heuer, H., Szczepanowski, R., Schneiker, S., Puhler, A., Top, E. M. and Schluter, A. (2004) The complete sequences of plasmids pB2 and pB3 provide evidence for a recent ancestor of the IncP-1 beta group without any accessory genes. Microbiology (Reading, Engl.), 150, 3591–3599.

Horemans, B., Bers, K., Ruiz Romero, E., Pose Juan, E., Dunon, V., De Mot, R. and Springael, D. (2016) Functional redundancy of linuron degradation in microbial communities in agricultural soil and biopurification systems. Appl. Environ. Microbiol., 82, 2843–2853.

Horemans, B., Hofkens, J., Smolders, E. and Springael, D. (2014) Biofilm formation of a bacterial consortium on linuron at micropollutant concentrations in continuous flow chambers and the impact of dissolved organic matter. FEMS Microbiol. Ecol., 88, 184–194.

Huerta-Cepas, J., Forslund, K., Coelho, L. P., Szklarczyk, D., Jensen, L. J., von Mering, C. and Bork, P. (2017) Fast genome-wide functional annotation through orthology assignment by eggNOG-Mapper. Mol. Biol. Evol., 34, 2115–2122.

Ito, H. and Iizuka, H. (1971) Taxonomic studies on a radio-resistant pseudomonas. Agricultural and Biological Chemistry, 35, 1566–1571.

Jain, A. and Srivastava, P. (2013) Broad host range plasmids. FEMS Microbiol. Lett., 348, 87–96.

Jechalke, S., Dealtry, S., Smalla, K. and Heuer, H. (2013) Quantification of IncP-1 plasmid prevalence in environmental samples. Appl. Environ. Microbiol., 79, 1410–1413.

Koboldt, D. C., Zhang, Q., Larson, D. E., Shen, D., McLellan, M. D., Lin, L., Miller, C. A., Mardis, E. R., Ding, L. and Wilson, R. K. (2012) VarScan 2: somatic mutation and copy number alteration discovery in cancer by exome sequencing. Genome Res., 22, 568–576.

Lagesen, K., Hallin, P., R?dland, E. A., Staerfeldt, H. H., Rognes, T. and Ussery, D. W. (2007) RNAmmer: consistent and rapid annotation of ribosomal RNA genes. Nucleic Acids Res., 35, 3100–3108.

Larsen, M. H., Biermann, K., Tandberg, S., Hsu, T. and Jacobs, W. R. (2007) Genetic manipulation of *Mycobacteriumtuberculosis*. Curr Protoc Microbiol, Chapter 10, Unit 10A.2.

Lefort, V., Desper, R. and Gascuel, O. (2015) FastME 2.0: A Comprehensive, accurate, and fast distance-based phylogeny inference program. Mol. Biol. Evol., 32, 2798–2800.

Letunic, I. and Bork, P. (2019) Interactive Tree Of Life (iTOL) v4: recent updates and new developments. Nucleic Acids Res., 47, W256–W259.

Li, H. and Durbin, R. (2009) Fast and accurate short read alignment with Burrows-Wheeler transform. Bioinformatics, 25, 1754–1760.

Li, H., Handsaker, B., Wysoker, A., Fennell, T., Ruan, J., Homer, N., Marth, G., Abecasis, G. and Durbin, R. (2009) The Sequence Alignment/Map format and SAMtools. Bioinformatics, 25, 2078–2079.

Li, X., Top, E. M., Wang, Y., Brown, C. J., Yao, F., Yang, S., Jiang, Y. and Li, H. (2014) The broad-host-range plasmid pSFA231 isolated from petroleum-contaminated sediment represents a new member of the PromA plasmid family. Front Microbiol, 5, 777.

Li, X., Wang, Y., Brown, C. J., Yao, F., Jiang, Y., Top, E. M. and Li, H. (2016) Diversification of broad host range plasmids correlates with the presence of antibiotic resistance genes. FEMS Microbiol. Ecol., 92.

Liu, Y., Lai, Q., Goker, M., Meier-Kolthoff, J. P., Wang, M., Sun, Y., Wang, L. and Shao, Z. (2015) Genomic insights into the taxonomic status of the *Bacillus cereus* group. Sci Rep, 5, 14082.

Lopez-Gutierrez, J. C., Henry, S., Hallet, S., Martin-Laurent, F., Catroux, G. and Philippot, L. (2004) Quantification of a novel group of nitrate-reducing bacteria in the environment by real-time PCR. J. Microbiol. Methods, 57, 399–407.

Lê, S., Josse, J. and Husson, F. (2008) Factominer: An R package for multivariate analysis. J Stat Softw, 25, 1–18.

Madeira, F., Park, Y. M., Lee, J., Buso, N., Gur, T., Madhusoodanan, N., Basutkar, P., Tivey, A. R. N., Potter, S. C., Finn, R. D. and Lopez, R. (2019) The EMBL-EBI search and sequence analysis tools APIs in 2019. Nucleic Acids Res., 47, W636–W641.

Martinez, B., Tomkins, J., Wackett, L. P., Wing, R. and Sadowsky, M. J. (2001) Complete nucleotide sequence and organization of the atrazine catabolic plasmid pADP-1 from *Pseudomonas* sp. strain ADP. J. Bacteriol., 183, 5684–5697.

Martini, M. C., Albicoro, F. J., Nour, E., Schluter, A., van Elsas, J. D., Springael, D., Smalla, K., Pistorio, M., Lagares, A. and Del Papa, M. F. (2015) Characterization of a collection of plasmid-containing bacteria isolated from an on-farm biopurification system used for pesticide removal. Plasmid, 80, 16–23.

Martini, M. C., Wibberg, D., Lozano, M., Torres Tejerizo, G., Albicoro, F. J., Jaenicke, S., van Elsas, J. D., Petroni, A., Garcillan-Barcia, M. P., de la Cruz, F., Schluter, A., Puhler, A., Pistorio, M., Lagares, A. and Del Papa, M. F. (2016) Genomics of high molecular weight plasmids isolated from an on-farm biopurification system. Sci Rep, 6, 28284.

Meier-Kolthoff, J. P., Auch, A. F., Klenk, H. P. and Goker, M. (2013) Genome sequence-based species delimitation with confidence intervals and improved distance functions. BMC Bioinformatics, 14, 60.

Meier-Kolthoff, J. P. and Goker, M. (2019) TYGS is an automated high-throughput platform for state-of-the-art genome-based taxonomy. Nat Commun, 10, 2182.

Mela, F., Fritsche, K., Boersma, H., van Elsas, J. D., Bartels, D., Meyer, F., de Boer, W., van Veen, J. A. and Leveau, J. H. (2008) Comparative genomics of the pIPO2/pSB102 family of environmental plasmids: sequence, evolution, and ecology of pTer331 isolated from Collimonas fungivorans Ter331. FEMS Microbiol. Ecol., 66, 45–62.

Nies, D. H. (1995) The cobalt, zinc, and cadmium efflux system CzcABC from *Alcaligenes eutrophus* functions as a cation-proton antiporter in *Escherichia coli*. J. Bacteriol., 177, 2707–2712.

Parks, D. (2018) Comparem. https://github.com/dparks1134/CompareM.

R Core Team (2019) R: A Language and Environment for Statistical Computing. R Foundation for Statistical Computing, Vienna, Austria. URL: https://www.R-project.org/.

San Millan, A. and MacLean, R. C. (2017) Fitness Costs of Plasmids: a Limit to Plasmid Transmission. Microbiol Spectr, 5.

Schluter, A., Szczepanowski, R., Puhler, A. and Top, E. M. (2007) Genomics of IncP-1 antibiotic resistance plasmids isolated from wastewater treatment plants provides evidence for a widely accessible drug resistance gene pool. FEMS Microbiol. Rev., 31, 449–477.

Schneiker, S., Keller, M., Droge, M., Lanka, E., Puhler, A. and Selbitschka, W. (2001) The genetic organization and evolution of the broad host range mercury resistance plasmid pSB102 isolated from a microbial population residing in the rhizosphere of alfalfa. Nucleic Acids Res., 29, 5169–5181.

Seemann, T. (2014) Prokka: rapid prokaryotic genome annotation. Bioinformatics, 30, 2068–2069.

Sen, D., Van der Auwera, G. A., Rogers, L. M., Thomas, C. M., Brown, C. J. and Top, E. M. (2011) Broad-host-range plasmids from agricultural soils have IncP-1 backbones with diverse accessory genes. Appl. Environ. Microbiol., 77, 7975–7983.

Sota, M., Tsuda, M., Yano, H., Suzuki, H., Forney, L. J. and Top, E. M. (2007) Region-specific insertion of transposons in combination with selection for high plasmid transferability and stability accounts for the structural similarity of IncP-1 plasmids. J. Bacteriol., 189, 3091–3098.

Sota, M., Yano, H., Nagata, Y., Ohtsubo, Y., Genka, H., Anbutsu, H., Kawasaki, H. and Tsuda, M. (2006) Functional analysis of unique class II insertion sequence IS*1071*. Appl. Environ. Microbiol., 72, 291–297.

Stamatakis, A. (2014) RAxML version 8: a tool for phylogenetic analysis and post-analysis of large phylogenies. Bioinformatics, 30, 1312–1313.

Szpirer, C., Top, E., Couturier, M. and Mergeay, M. (1999) Retrotransfer or gene capture: a feature of conjugative plasmids, with ecological and evolutionary significance. Microbiology (Reading, Engl.), 145 (Pt 12), 3321–3329.

Tauch, A., Schneiker, S., Selbitschka, W., Puhler, A., van Overbeek, L. S., Smalla, K., Thomas, C. M., Bailey, M. J., Forney, L. J., Weightman, A., Ceglowski, P., Pembroke, T., Tietze, E., Schroder, G., Lanka, E. and van Elsas, J. D. (2002) The complete nucleotide sequence and environmental distribution of the cryptic, conjugative, broad-host-range plasmid pIPO2 isolated from bacteria of the wheat rhizosphere. Microbiology (Reading, Engl.), 148, 1637–1653.

Thomas, C. M., Thomson, N. R., Cerdeno-Tarraga, A. M., Brown, C. J., Top, E. M. and Frost, L. S. (2017) Annotation of plasmid genes. Plasmid, 91, 61–67.

Vedler, E., Vahter, M. and Heinaru, A. (2004) The completely sequenced plasmid pEST4011 contains a novel IncP1 backbone and a catabolic transposon harboring *tfd* genes for 2,4-dichlorophenoxyacetic acid degradation. J. Bacteriol., 186, 7161–7174.

Thorsted, P. B., Shah, D. S., Macartney, D., Kostelidou, K. and Thomas, C. M. (1996) Conservation of the genetic switch between replication and transfer genes of IncP plasmids but divergence of the replication functions which are major host-range determinants. Plasmid, 36, 95–111.

Yan, Z., Zhang, Y., Wu, H., Yang, M., Zhang, H., Hao, Z. and Jiang, H. (2017) Isolation and characterization of a bacterial strain *Hydrogenophaga* sp. pyr1 for anaerobic pyrene and benzo[a]pyrene biodegradation. RSC Adv., 7, 46690–46698.

Yanagiya, K., Maejima, Y., Nakata, H., Tokuda, M., Moriuchi, R., Dohra, H., Inoue, K., Ohkuma, M., Kimbara, K. and Shintani, M. (2018) Novel self-transmissible and broad-host-range plasmids exogenously captured from anaerobic granules or cow manure. Front Microbiol, 9, 2602.

Zhang, M., Visser, S., Pereira e Silva, M. C. and van Elsas, J. D. (2015) IncP-1 and PromA group plasmids are major providers of horizontal gene transfer capacities across bacteria in the mycosphere of different soil fungi. Microb. Ecol., 69, 169–179.

Öztürk, B., Werner, J., Meier-Kolthoff, J. P., Bunk, Spröer, C. and Springael, D. (2019) Comparative genomics unravels mechanisms of genetic adaptation for the catabolism of the phenylurea herbicide linuron in *Variovorax*. bioRxiv 759100; doi: https://doi.org/10.1101/759100

